# Detection of Long COVID Microclots using Pulsed Speckle Contrast Optical Spectroscopy

**DOI:** 10.1101/2025.08.06.669007

**Authors:** Reza Rasouli, Brad Hartl, Soren D. Konecky

**Affiliations:** Openwater, San Francisco, California, United States

## Abstract

Circulating microclots are increasingly linked to long COVID as well as its persistent symptoms such as fatigue, cognitive deficits, and cardiovascular complications. These conditions can become debilitating or even life-threatening, which create an urgent need for rapid and reliable detection and monitoring tools. In this study we investigate pulsed speckle contrast optical spectroscopy (p-SCOS) as a non-invasive and label-free method to detect microclots in biofluids. Microclots at four concentrations (21k, 91k, 400k, and 1.7M microclots/mL), representing levels from healthy individuals to acute coagulopathic states, were generated using a freeze-thaw method. We measured speckle contrast under flowing conditions in a custom-made flow phantom. In phosphate-buffered saline (PBS) and plasma, increasing microclot concentration consistently led to measurable decreases in speckle contrast. The measurement differentiated between low and high clot burdens in transparent media which highlights its potential for microclot monitoring. In comparison, no detectable changes were observed in whole blood, likely due to dominant scattering from red blood cells masking microclot effects. Overall, our findings demonstrate the feasibility of p-SCOS as a rapid and label-free tool for microclot detection and monitoring in transparent biofluids.

## Introduction

Amyloid microclots are fibrin(ogen)-based aggregates ranging from 1 to 200 µm in diameter. Unlike normal fibrin structures, these clots exhibit β-sheet-rich conformations that render them resistant to fibrinolysis.^1–3^ Kell and colleagues have shown that amyloid fibrin microclots persist in circulation and bind to inflammatory mediators which in turn create a pro-thrombotic and pro-inflammatory environment.^4–6^ When deposited in capillaries and small arterioles, these clots can obstruct microvascular flow and limit oxygen delivery which in turn leads to local hypoxic conditions. These microclots appear to contribute to the pathophysiology of long COVID, where persistent symptoms such as fatigue, cognitive impairment, and cardiovascular dysfunction occur.^2,7–10^ Microclots have been consistently identified in the plasma of long COVID patients and suggest a sustained imbalance in coagulation and clearance.^6,11,12^ Similar fibrin amyloid microclots have been reported in other inflammatory and metabolic diseases, including Alzheimer’s, Parkinson’s, and type 2 diabetes, where they may impair microvascular perfusion and promote chronic inflammation.^13,14^ In these contexts, microclots may act as upstream drivers of inflammation and tissue stress, through a shared mechanism of vascular occlusion and impaired perfusion.

Despite the clinical relevance of microclots, quantitative methods for their detection remain limited. Long COVID is currently diagnosed based on clinical history and self-reported symptoms, without an objective biomarker for microclot burden.^9,11^ Laboratory approaches have relied on staining patient plasma samples with amyloid-binding dyes (e.g., Thioflavin T) followed by fluorescence microscopy.^15^ While this method has successfully revealed elevated microclot loads, it is invasive, labor-intensive, and impractical for large-scale or point-of-care use. It requires blood collection, specialized reagents, fluorescence microscopic imaging equipment, and expert interpretation. This diagnostic gap limits both clinical diagnosis and routine monitoring for emerging therapies aimed at resolving microclots. Given the dynamic nature of microvascular pathology in systemic coagulopathies, there is a need for a method that enables reliable, label-free monitoring over time. An approach that supports non-invasive measurements could substantially advance both clinical management and research into microclot-associated conditions.

Optical methods offer a promising alternative for rapid and label-free microclot detection. In particular, pulsed Speckle Contrast Optical Spectroscopy (p-SCOS),^16–18^ is a technique sensitive to motion and microstructural changes in scattering media which renders it a promising approach for monitoring blood microclots. In p-SCOS, highly coherent pulsed laser light is delivered to the sample (e.g., blood or plasma), and the returned multiply scattered light produces a random interference pattern of bright and dark speckles when resolved on an imaging sensor. The temporal fluctuations of this speckle pattern depend on the movement of scatterers (cells or particles) within the sample. Faster or more abundant motion blurs the speckle pattern and decreases the measured contrast, whereas static or slowly moving scatterers result in higher contrast.^19^ Thus, changes in blood flow dynamics or the presence of particulate microstructures like clots can be detected as changes in speckle contrast.

A major advantage of p-SCOS is its label-free nature as it requires no external contrast agents or chemical processing, enabling real-time assessment over large sample volumes.^20^ Speckle contrast-based methods have been applied in biomedical research for non-invasive blood flow monitoring, including in cerebral perfusion, wound assessment, and stroke detection.^19,21–23^ These features suggest that p-SCOS could be adapted to detect microclots in biofluids.

In this study, we aim to assess the feasibility and limitations of p-SCOS as a non-invasive method for detecting and quantifying microclot burden under flowing conditions. We hypothesize that microclots will induce measurable changes in speckle contrast of biofluids. We introduce known concentrations of microclots ranging from 21k (a measured value of healthy donors), to 1.7 M microclots/mL (to represent acute coagulopathic into samples)^11^ and measure the speckle contrast values to determine the sensitivity of p-SCOS to incremental microclot burdens. We compared the performance of p-SCOS to detect microclots in common biofluids such as PBS, plasma, and in whole blood.

## Materials and Methods

### Materials

Porcine plasma was obtained from Innovative Research, Inc. (USA) and stored at −20 °C until use. Plasma aliquots (15 mL) were used to prepare microclots.

### Sample Preparation

Amyloid microclots were prepared by subjecting 15 mL aliquots of porcine plasma (Innovative Research, USA) to 12 freeze-thaw cycles (−20 °C to 37 °C, 2 hours per cycle), followed by incubation at 37 °C.^24^ Microclot suspensions were prepared at four concentrations: 21,000, 91,000, 400,000, and 1.7 million microclots/mL. For whole blood experiments, the 1.7M/mL microclot suspension was diluted into fresh whole blood to approximate physiological hematocrit while maintaining high clot burden. Plasma and PBS were used as more transparent media for contrast testing. Thioflavin T (Sigma-Aldrich, USA) was used for amyloid staining in endpoint assays.

### p-SCOS Setup

Speckle contrast was performed using the Openwater Open-Motion 2.0 system described in greater detail previously.^17^ Illumination was provided by a pulsed 785 nm laser (250 μs pulse width, 40 Hz, 400 μJ/pulse). The returned light was collected via a 1.5 mm aperture and a 5-megapixel CMOS camera (HM5530, Himax Technologies, Taiwan), located 12 mm from the laser output (see Figure 1). Custom python scripts as described previously were used to calculate speckle contrast.^17^

**Figure 1.**
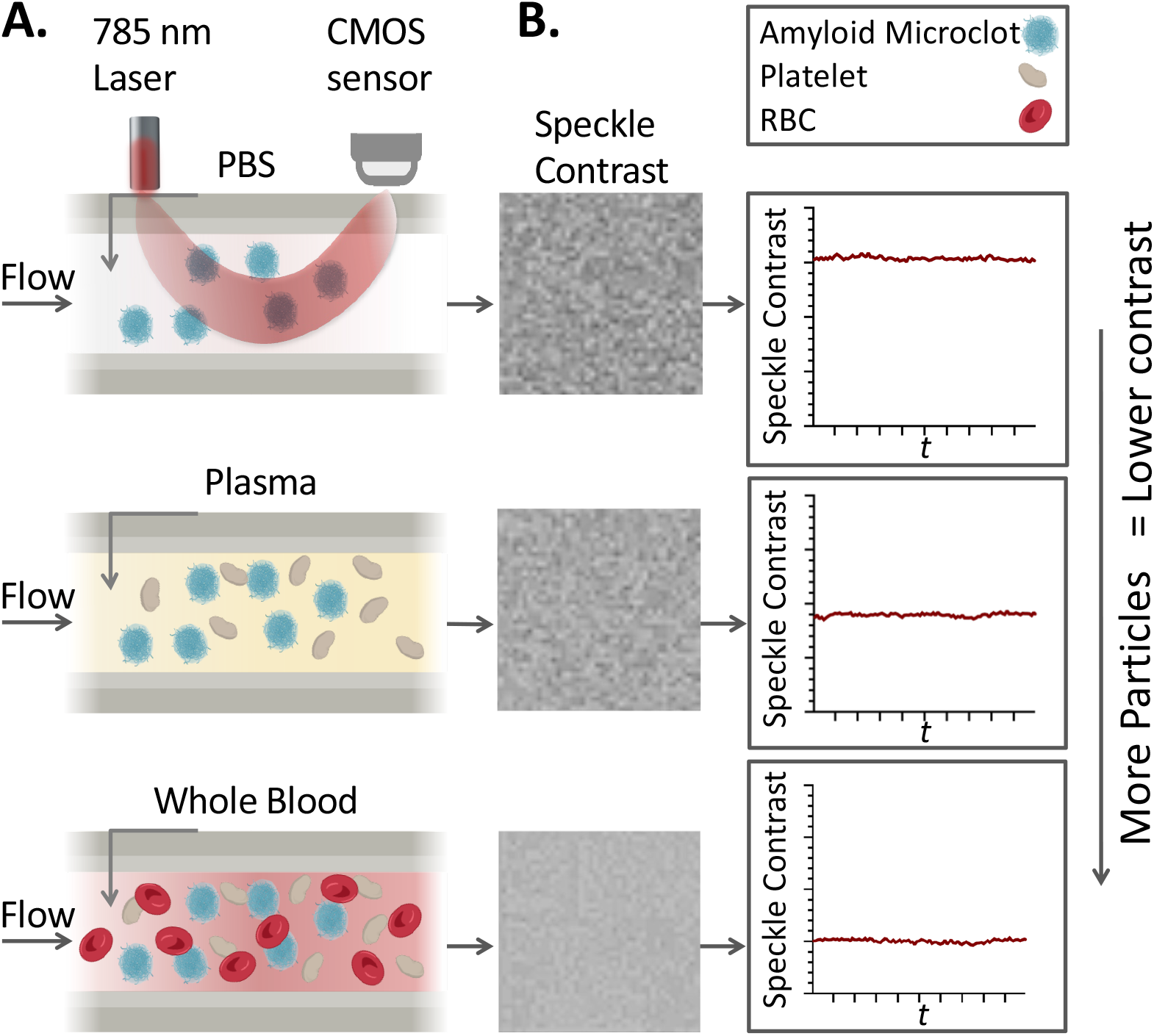
Overview of the speckle contrast detection system for microclot assessment under flow. **A.** Schematic of the illumination and detection within the optical phantom for detection in PBS, plasma, and whole blood, each containing varying numbers of amyloid microclots, platelets, and red blood cells. **B**. Representative speckle images corresponding to each sample type and speckle contrast traces over time where higher contrast indicates a greater number of scattering particles.

### Phantom Design and Fabrication

Biofluids were flowed through a highly scattering optical phantom with optical properties similar to tissue. The phantom contained a channel with a center depth of 4 mm. The flow phantom was cast from optically clear urethane resin (Smooth-On Crystal Clear 200) with embedded scattering and absorbing agents (1.0 g/L ~325 mesh TiO_2_ and 0.020 g/L carbon black). 3D-printed molds were used to define channel geometry (3.175 mm diameter) and components were mixed and degassed before curing. A syringe pump was used to flow the biofluids through the channel at 20 mL/min (equivalent to 4.2 mm/s).

### Statistical Analysis

All experiments were repeated in at least two independent batches. Mean speckle contrast values were compared across concentrations and sample types using ANOVA with Tukey’s post-hoc testing (significance threshold p = 0.05). For each batch, technical replicates consisted of multiple data points (n = 500-700 per repetition). To avoid underestimating variability and inflating significance, pooled standard deviation formula was calculated by:

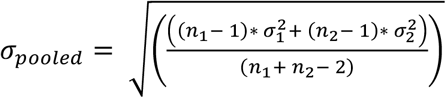 where σ_1_ and σ_2_ are the standard deviations of each repetition, and n_1_ and n_2_ are the number of measurements per repetition. The resulting pooled standard deviation was used for error bars Error bars represent the pooled standard deviation unless otherwise noted. Significance levels are denoted as *p* < 0.05 (\**), p < 0*.*01 (**), p < 0*.*001 (\*\*\**), and *p < 0*.*0001 (\*\*\*\**).

## Results

We evaluated the performance of speckle contrast microclot detection in three media of increasing optical complexity. First, transparent PBS provided a near-optimal scattering baseline. Next, plasma added physiologic turbidity from soluble proteins while remaining free of blood cells. Finally, whole blood which clinically is the most relevant, yet optically the most challenging, introduced dense red- and white-blood cell scattering. This tiered approach allowed us to **(a)** establish the maximum contrast separation amongst groups under ideal conditions, **(b)** assess detection sensitivity in a biologically meaningful fluid, and **(c)** define the limits imposed by dense cellular scattering.

Microclot suspensions at four representative concentrations were studied: 21k clots mL^−1^ (median in healthy donors), 91k clots mL^−1^ (~2× the long-COVID median), 400k clots mL^−1^, and 1700k clots mL^−1^. Two independent microclot batches (termed B1 and B2) were introduced into each medium and analyzed under identical flow conditions.

### Microclot Detection in PBS

Under flow, speckle contrast of microclots in PBS showed a clear, concentration-dependent response to microclot burden. **Figure 2** shows time-averaged contrast values from two independent experimental batches (B1, B2). At the lowest concentration (21k clots mL^−1^, healthy baseline), contrast was highest (0.31 for B1 and 0.32 for B2). Increasing the microclot count to 91k mL^−1^ and 400k mL^−1^ produced stepwise declines, while the highest load (1.7M mL^−1^) resulted in the lowest contrast (0.21 and 0.22). One-way ANOVA confirmed a significant effect of concentration on contrast between groups (p < 0.05). No plateau or reversal was observed even at the highest density, suggesting clots remained in continuous flow and did not cause measurable flow stasis in the interrogated region. Reproducibility between acquisition was high and both batches showed comparable drop in contrast across concentrations.

**Figure 2.**
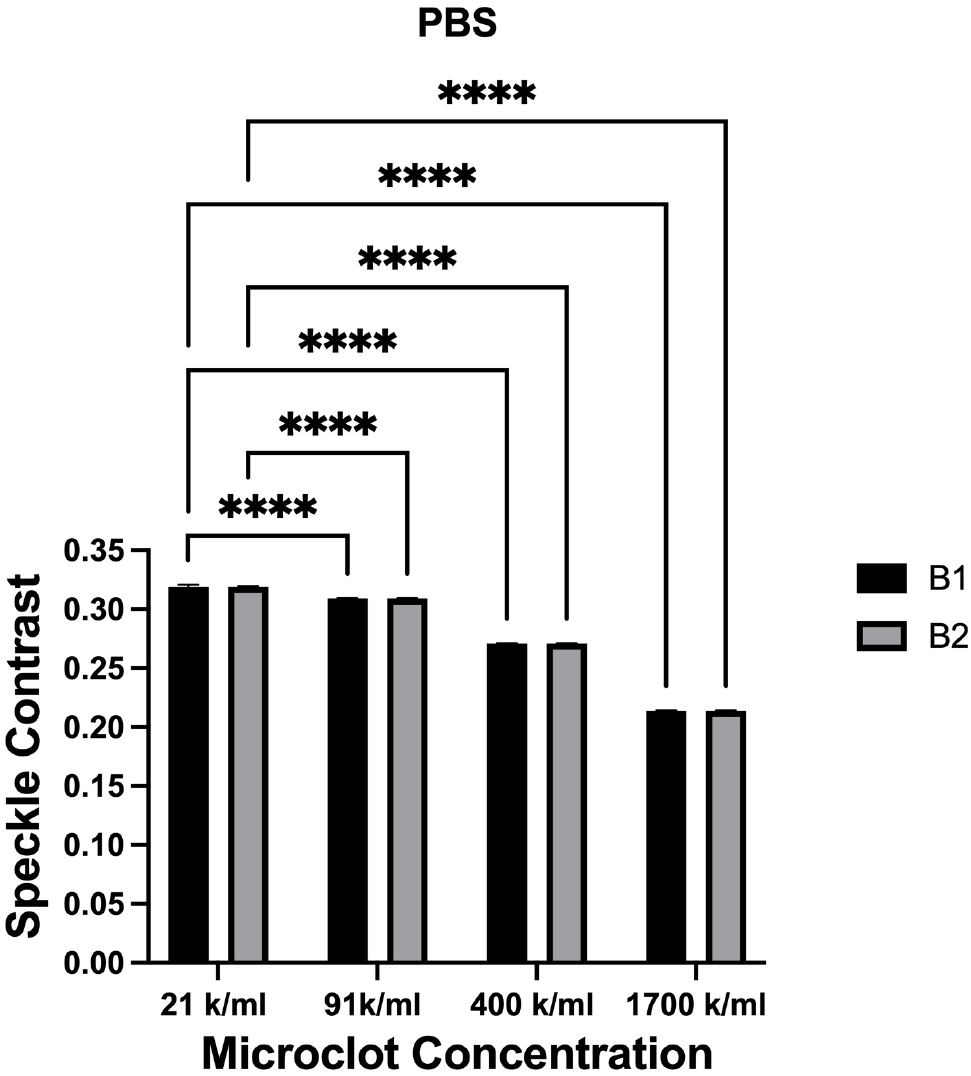
Averaged contrast values across all concentrations (21 k/mL, 91 k/mL, 400 k/mL, and 1.7 M/mL) in PBS for both experimental batches. Error bars represent pooled standard deviation from replicate measurements. Contrast consistently decreased with increasing clot concentration, demonstrating the system’s ability to distinguish clot burden under idealized optical conditions.

### Microclot Detection in Plasma

In plasma, baseline contrast values were lower than in PBS which reflect the medium’s increased number of scatterers. However, the concentration-dependent trend seen in PBS was repeated in plasma as well. **Figure 3** shows that at 21k clots mL^−1^, contrast averaged 0.26 for B1 and 0.30 for B2, dropping to 0.24 and 0.28 respectively at 400k mL^−1^, and at 1.7M mL^−1^ it fell to 0.21 for B1 and 0.25 for B2. The overall dynamic range (~ 0.05 absolute) represents a ~20 % reduction from baseline. ANOVA indicated a significant concentration effect (p < 0.05) in both batches. However, the significance between 21k and 91k was inconsistent between batches. Although batch 2 yielded slightly higher absolute contrast than batch 1 (likely due to intrinsic differences in plasma content)^25–27^, the monotonic decrease with increasing clot load was preserved. The smaller contrast span relative to PBS is consistent with greater baseline scattering that partially masks clot-induced speckle decorrelation, but the method nonetheless detects microclot increments in this biologically relevant fluid.

**Figure 3.**
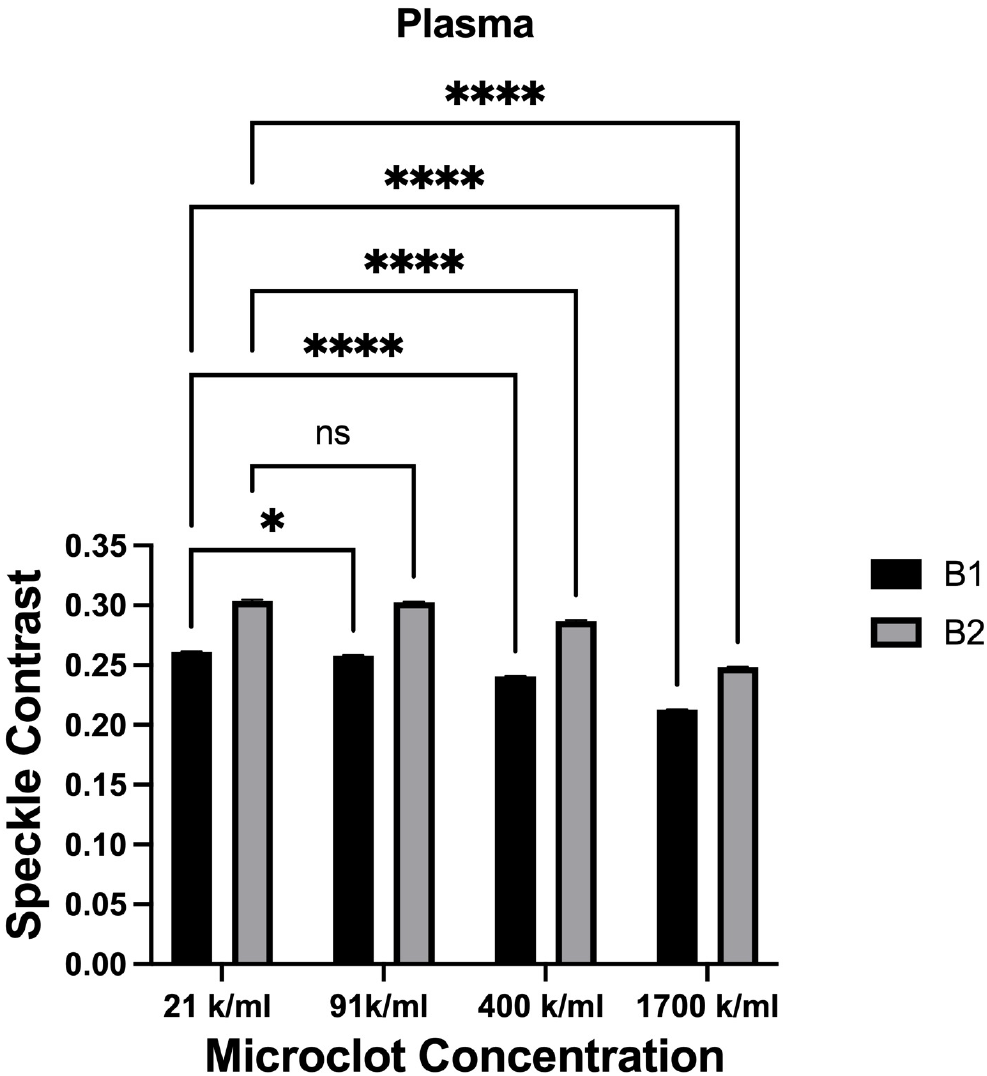
Mean contrast values for all concentrations in plasma, averaged over both acquisitions. Error bars represent pooled standard deviation from replicate measurements. The results show a concentration-dependent decrease in contrast, indicating that the system is effective for detecting high microclots shifts even in more complex but with low cell density biological fluids.

### Microclot Detection in Whole Blood

Whole-blood measurements revealed reduced contrast modulation across the microclot concentration range. **Figure 4** shows the mean contrast values clustered narrowly around 0.20 at 21k clots mL^−1^ and remained in the same range for all concentrations. ANOVA found no statistically significant effect of concentration. Replicate batches confirmed the outcome. No contrast increase was observed that would suggest flow obstruction and cellular motions seem to govern speckle decorrelation regardless of additional clots. These findings highlight that in highly scattering and cell-dense media, speckle contrast measurements might not reliably be able to resolve the small contribution of physiologically relevant microclot loads.

**Figure 4.**
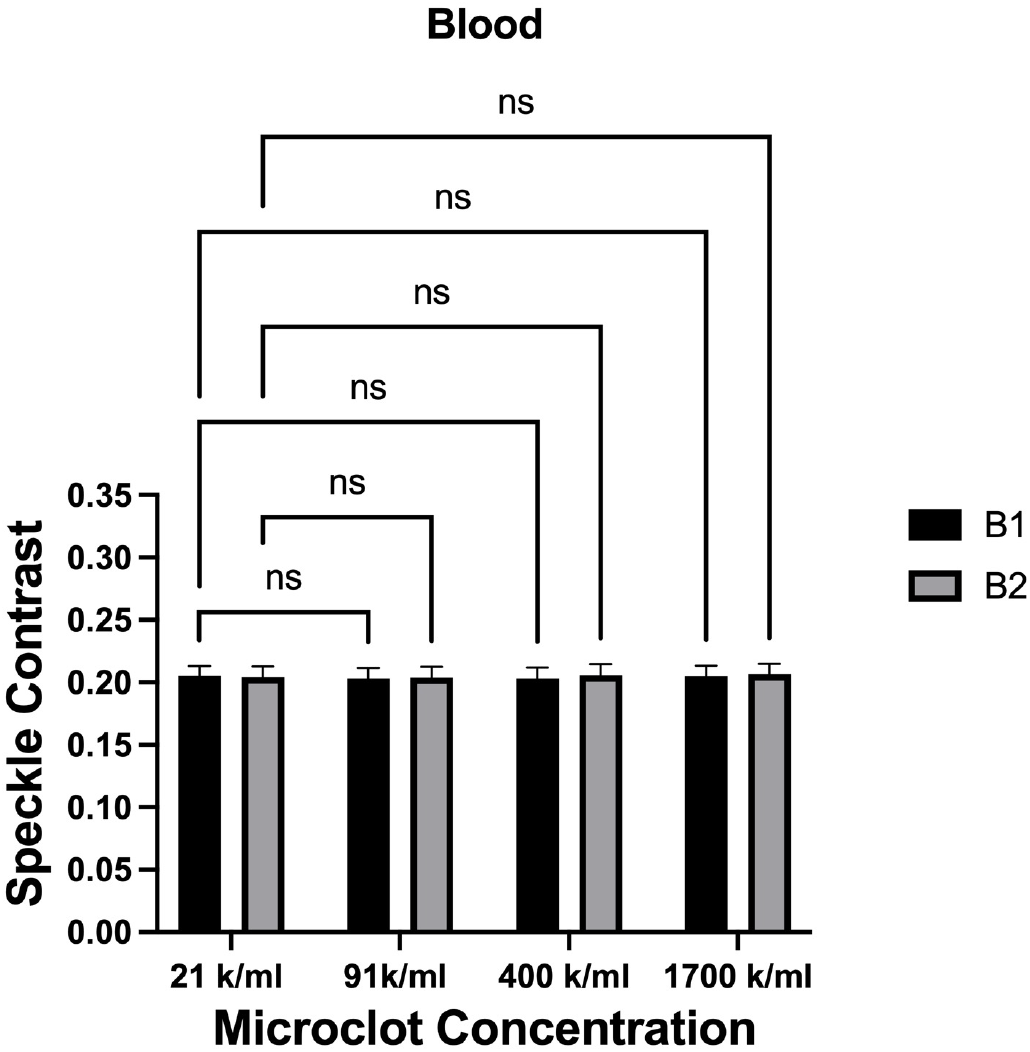
Averaged data for whole blood across all concentrations and batches. Error bars represent pooled standard deviation. No statistically significant differences were detected between groups. The scattering from whole blood cell populations likely obscures microclot-induced changes in speckle contrast.

## Conclusion

Developing a rapid, label-free microclot detection tool would address a critical unmet need in the management of long COVID and related disorders. An optical, speckle contrast-based microclot assay could potentially be used to monitor disease progression or response to treatments in patients with post-COVID syndrome and other thrombo-inflammatory conditions, systemic coagulation disorders, or even metabolic and neurodegenerative diseases. This proof-of-concept study demonstrates the potential of p-SCOS for detecting clot-related pathologies in real time. Across batches, speckle contrast reliably distinguished four microclot levels in optically clear (PBS) and moderately scattering (plasma) fluids, with statistically significant, monotonic contrast differences as clot counts increased. The technique’s sensitivity stems from the inverse relationship between speckle contrast and the combined density and speed of scatterers. In whole blood, however, the microclot signal was obscured by the presence of red and white blood cells. These results suggest that clinical translation may require either plasma-based measurements, or additional signal- and/or sample-processing approaches to extract microclot signatures from whole-blood speckle contrast data. The present study showed the feasibility and limitations of p-SCOS technologies for non-invasive microclot screening in long COVID and other coagulopathic conditions, enabling earlier intervention and better tracking of patient health in these challenging syndromes.

## Data availability statement

The original contributions presented in the study are included in the article/supplementary material, further inquiries can be directed to the corresponding author.

